# Reactive oxygen species and evaluation of the quality of carcasses and beef meat from cold storage slaughterhouses located in the Federal District and Surroundings (Brazil)

**DOI:** 10.1101/490342

**Authors:** Thaís Rezende Leite, Carla Magalini Zago de Sousa, Paulo Eduardo Narciso de Souza, Fabiane Hiratsuka Veiga de Souza, Vítor Salvador Picão Gonçalves, Ana Lourdes Arrais de Alenca, Aline Mondini Calil Racanicci, Angela Patrícia Santana

**Author notes:** These authors contributed equally to this work (Thaís Rezende Leite, Carla Magalini Zago de Sousa, Angela Patrícia Santana). These authors contributed equally to this work (Paulo Eduardo Narciso de Souza, Fabiane Hiratsuka Veiga de Souza). These authors contributed equally to this work (Vítor Salvador Picão Gonçalves, Ana Lourdes Arrais de Alenca). This author contributed to this work (Aline Mondini Calil Racanicci).

## Abstract

The aim of this study was to detect reactive oxygen species and evaluate the quality of carcasses and beef meat from cold storage slaughterhouses located in the Federal District and Surroundings. Information was obtained on the gender, breed and age of each animal, as well as the distance travelled (km) and time (h) from the property to the cold storage slaughterhouse. Data and samples were obtained from a total of 33 animals and their respective carcasses. Fragments of the extensor carpi radialis muscle were extracted from all the carcasses to analyze the presence of reactive oxygen species (ROS), and- samples of the *Longissimus dorsi* muscle to perform the 24h post-mortem pH, colorimetric test, cooking loss and drip loss assessment and to measure the shear force. The presence of hematomas was detected in 28 carcasses, where the tail and croup (17/33) and flank (17/33) regions were the most affected. The electron paramagnetic resonance indicated an average of 52.59 ROS/g in the analyzed pieces. The meat quality tests indicated averages of: 5.8 for the 24h postmortem pH, L*29.34, a* 2.52 and b* 1.31 in calorimetry, 2.30 kg/f for the shearing force, 11.75% of cooking losses and 1.88% of drip losses. The statistical analyses demonstrated a tendency- to positive correlation between the presence of hematomas with and the amount of ROS, and between the presence of hematomas and pH value. Furthermore, statistically, female gender was one of the influential factors on the tenderness value. According to the results, it was concluded that the meat evaluated in this study meets the desirable quality parameters and it was possible to detect reactive oxygen species in the samples of muscle tissue.

## Introduction

Brazil’s livestock farming stands out as an important sector of the national economy and in the year 2016 it represented approximately 24% of the Gross Domestic Product [1]. In 2017 Brazil kept its spotlight in the international market as the world’s greatest beef meat exporter [1].

To maintain its market competitiveness, the country has invested in the productivity and quality of the final product offered to the consumer [2]. In beef, tenderness, flavor, juiciness and color are desired as determining characteristics for the purchase decision, which are susceptible to changes due to intrinsic factors such as the gender, age and breed, and extrinsic factors such as nutrition and stress during handling [3].

Studies on meat quality often correlate these parameters when evaluating the characteristics of the meat produced by crossbreeding and the comparison of age and diet types [4,5,6,7], where the main analyzed characteristics are the pH, colorimetry and shearing force [8,9,10] as well as water retention capacity when measured by evaluating cooking and drip losses [4].

Still related to the meat quality is the pre-slaughter handling performed by the farms and industries of meat, which can generate stress, contusions and hematomas in the animal, interfering with meat quality at the end of the process and causing economic losses to the producer and the industry [11, 12; 13].

Electron paramagnetic resonance spectroscopy (EPR) is a technique that has been applied to foods, making it possible to detect free radicals at an intracellular and extracellular level [14], thus allowing the correlation of the presence of free radicals with the quality of the final product [15]. Chen et al. [16] used the EPR technique to evaluate the oxidative stress mechanism in broiler chickens. The authors confirmed that oxidative stress induced by H2O2 had a negative effect on relative muscle weight, the significantly higher ROS formation in the muscle show lower quality meat, with lower pH24h value, higher shearing force and greater drip losses [16]. Another study applied EPR to measure the meat quality of pigs shortly after slaughter [17]. The authors observed that oxidation occurred differently in the different muscle tissues and that this process can reduce meat quality, changing the flavor, coloration, drip and cooking losses.

To this moment no scientific findings were found to correlate the detection of reactive oxygen species with the beef meat quality in combination with the pre-slaughter handling of the livestock. In the region of the Federal District and Surroundings there is no information on the characteristics of beef meat quality meant for sale in terms of the pH, tenderness, cooking and drip losses, as well as the characterization of the pre-slaughter handling and their consequences in the conversion of muscle into meat in the daily routine of the regional cold storage industry.

Considering the importance of the behavior of the organoleptic characteristics of beef meat, as well as the pre-slaughter handling, this project aimed to evaluate the 24h postmortem pH and coloration, the drip and cooking liquid losses and the shearing force in beef meat from cold storage slaughterhouses located in the Federal District and Surroundings, and to detect reactive oxygen species in muscle tissues by electron paramagnetic resonance.

## Materials and methods

### Origin of the samples

The collection of data and samples was performed in cold storage beef slaughterhouses located in the Federal District and Surroundings. All slaughterhouses were certified by an official inspection service such as SIF or SISBL and the animals were subject to the norms of Humane Slaughter outlined in the Normative Instruction no. 3, of January 17th, 2000, according to the Ministry of Agriculture, Livestock and Supply [18]. Four collections were performed between April and September, 2017 and 33 beef carcasses were obtained in total.

During the pre-slaughter handling in the cold storage slaughterhouses, information was obtained on the travel time (h) and distance (km) from the location of the rural property from which the animals originated, as well as the identification of the animals by batch, breed, age and gender. After the desensitization, bleeding and skinning procedures, pieces of approximately 3.0 cm of the extensor carpi radialis muscle were collected and frozen in liquid nitrogen for the subsequent quantification of reactive oxygen species. Next, the number and location of hematomas in the carcasses were evaluated, dividing them into regions proposed by the methodology by Cardoso et al., [19].

After the cooling of the 33 carcasses for 24h in the slaughterhouse refrigeration chamber, a piece of the *Longissimus dorsi* muscle of approximately 500g was removed, 2.5 cm thick, between the 10^th^ and 12^th^ rib for the meat quality analyses.

### Quantification of reactive oxygen species (ROS)

For the quantification of reactive oxygen species in the muscle tissue, the protocol recommended by Mrakic-Sposta et al. [20] was followed. The pieces of the extensor carpi radialis muscle immersed in liquid nitrogen were transported to the Laboratory of Biochemistry and Protein Chemistry at the University of Brasília to prepare the samples for the paramagnetic resonance spectroscopy analyses at the Laboratory of Electron Paramagnetic Resonance at the Institute of Physics.

### Analyses of beef meat quality

#### pH and colorimetric test

The pH was measured with a portable digital pHmeter (Testo® 205) equipped with an insertion electrode. The device was calibrated by immersion in buffer solution at pH 4.0, followed by one at pH 7.0, as suggested in the manufacturer’s manual. The samples of 24h post-mortem beef meat were exposed to atmospheric air for 30 minutes, followed by three readings at different points and the calculation of their arithmetic mean. A portable Chroma Meter Cr-400 (Minolta Camera Co., Ltda., Osaka, Japan) colorimeter was used. To determine the color, the CIELAB color space [21] was adopted, which applied the lightness (L*) coordinates varying from 0 (black) to 100 (white), shades of red (a*+) to green (a*−), and shades of yellow (b*+) to blue (b*−).

#### Cooking loss

To evaluate the weight loss during the cooking, the procedure recommended by AMSA [10] with adaptations was used as reference, where the cooled meat samples were weighed, then subjected to a cooking process in an industrial oven of brand Venancio®, until a temperature of 70°C was reached inside the pieces, thus obtaining the weight before cooking. The cooking temperature was monitored by a metal thermometer of brand Testo® 926 with needle probe model 06280030 for the control of the internal temperature of the sample. After reaching the internal temperature of 70°C, the samples were removed from the oven, cooled at room temperature and weighed again for the calculation of the cooking losses (Cooking loss = [(weight of raw steak - cooked weight) ÷ weight of raw steak] × 100). After this procedure, the samples were individually packaged in low-density polyethylene bags, properly identified and stored under refrigeration at 4°C for 24h for the subsequent performance of the shearing force test.

#### Shearing force

The protocol recommended by Wheeler et al. [9] was applied. After being stored at 4°C for 24h following the cooking test, the samples were prepared for the performance of the shearing force test. Three cylindrical pieces were collected from each meat sample, cut parallel to the direction of the muscle fiber using a perforator with a diameter of approximately 1.27 cm. These three sub samples underwent the shearing test using equipment of model WARNER-BRATZLER MEAT SHEAR® (G-R Manufacturing Co. Manhattan), 235 6X. The results were given in Kg/f and the mean between the three sub samples was calculated to represent the force used to cut each sample.

#### Drip loss

The drip loss was measured according to the procedure recommended by Honikel [22] with adaptations by Kim et al. [23].

A piece of approximately 50g was removed 24h post mortem from each beef meat sample from the refrigerated slaughterhouses. The sample was cleaned and the excess fat and external connective tissue were disposed for the weighing.

The samples were properly identified and individually placed in polyethylene nets. Theywere then suspended by a hook, wrapped by a polyethylene bag with no contact with the sample and stored under refrigeration at 4°C for 48 h. After this period the sample was weighed and the drip loss was calculated (Drip loss = [weight after the dripping ÷ weight before] × 100).

### Detection of the reactive oxygen species (ROS) in the muscle tissue

The samples of the pieces of the extensor carpi radialis muscle collected shortly after the bleeding and measuring approximately 3cm were frozen in liquid nitrogen and taken to the Laboratory of Biochemistry and Protein Chemistry of the University of Brasília, where they were subjected to freezing at −80°C until the preparation of the samples.

They were removed from the freezer and then cut with a scalpel, reduced to four parts of approximately 2×2×2 mm. Next, the tissues were washed thrice with a KHB solution (Krebs Hepes, Noxygen, Germany). After washing, 700 μL of CMH was added at a concentration of 200 μM (Noxygen®, Germany) and 50 UI/mL of sodium heparin (Hepamax-S®, Blausiegel Ind. Com. Ltda., São Paulo, Brazil) was added. Samples were incubated under agitation at 37°C for 60 minutes. Incubation was followed by the removal of 450 μL of the supernatant, which was transferred to a 1 mL syringe for immediate freezing in liquid nitrogen and subsequent storage at −80 °C until the reading by the electron paramagnetic resonance spectrometer.

The remaining pieces of muscle tissue in the *eppendorff* tubes were dehydrated in Speed Vac (Savant SC100) apparatus and weighed for the calculation of the reactive oxygen species in the muscle tissue.

### Electron paramagnetic resonance (EPR) measurements

The calibration of the electron paramagnetic resonance apparatus and the reactive oxygen species analysis were performed according to the protocol described by Gomes et al [24]. The measurements were performed at the Laboratory of Electron Paramagnetic Resonance of the Institute of Physics at the University of Brasília.

The X-band Bruker® EMX500 spectrometer apparatus was used (9.45 GHz), 2 mW of microwave power, 5 Gauss modulation field, modulation frequency of 100 kHz, scan width of 200 G, scan time of 10 s and 5 scans added in combination for each measurement.

The calibration curve was defined based on Berg et al. [25], where spectrometer measurements were performed with a 10 mM solution of the CP• (3-carboxy-proxyl) (Noxygen®, Germany) radical prepared in KHB buffer solution and diluted at 0.5, 10, 50, and 100 μM. Next, 450 μL of the calibration samples was transferred to a 1 mL polyethylene syringe (Descarpax®) and frozen in liquid nitrogen. These calibration samples generated the curve used for the quantification of the reactive oxygen species.

The frozen samples were removed from the syringes and placed individually in the Finger Dewar container (Noxygen®Germany), which was subsequently filled with liquid nitrogen. The Finger Dewar was then coupled to the spectrometer with each sample, still frozen, one at a time, and the detection of the reactive oxygen species was performed individually, which generated a curve corrected for the CP• quantity according to the calibration curve, obtaining the ROS quantification in the sample as the result.

### Statistical Analysis

The results were subjected to statistical analysis of the mean and standard deviation using the StataCorp. 2011 software. Stata: Release 12. Statistical of the Software. College Station, TX: StataCorp LP. The Shapiro-Wilk Test was applied to verify the normal distribution of the analyzed variables (pH and gender). The Kruskal-Wallis Test is a non-parametric test used to verify normality between the days of the collections. The Kolmogorov-Smirnov, also non-parametric, was used to compare hematoma and pH means, and hematoma and reactive oxygen species means.

## Results and discussion

The information obtained on the travel distances in (km) and travel time (h) from the property to the cold storage slaughterhouses, of the 33 cattle in this study, as well as the breed, gender and age are presented in Table 1.

**Table 1.** Data on the 33 cattle: travel distance (km) and travel time (h), breed, gender and age (months) obtained at the cold storage slaughterhouses in the 04 collections carried out from April to September, 2017.

| Collections | Distance (Km) | Travel time (h) | Rest time at the slaughterhouse (h) | Breed | Gender | Age in months |
| --- | --- | --- | --- | --- | --- | --- |
| 1 | 90 | 1h 20 m | 13h 30m | Crossbred | 9 males | 24 - 36 |
| 2 | 130 | 1h 40 m | 13h 30m | Crossbred | 4 males | 24 - 36 |
| 3 | 70 | 1h | 16h | Crossbred | 1 male<br>6 females | 24 - 36 |
| 4 | 140 | 2h | 13h | Crossbred | 13 females | 24 - 36 |
| Total |  |  |  |  | 33 animals |  |

The travel distances by road transportation observed in this study were on average 110 km and the mean travel time was 1h 50m. According to studies such as those of Joaquim [26]; Pereira et al., [27] and Batista De Deus, et al., [28], the travel time can influence the final quality, therefore, animals travelling long distances (greater than 330 km) can present higher final pH (pH>6) compared to the short distance travels [26, 27, 28].

The time from the arrival at the cold storage slaughterhouse to the moment of slaughter ranged from 13 to 16 hours, this rest time corroborates with RIISPOA [29], which requires rest and fasting of at least 6h at the slaughterhouse, with a possible extension to 24h, when the travel time does not exceed 2 hours for the cattle [29]. However, the latest version does not establish a minimum and maximum period of fasting and water diet [30].

Inevitably, longer travel distances involve longer fasting and water diet periods, which, for prolonged periods (longer than 16 hours) can affect meat quality, causing fatigue and stress to the animal that can lead to an elevated final pH and higher shearing force, and it can also lead to weight loss and dehydration in the cattle [31, 32].

The distances found in the present study have short travel times, not requiring long fasting periods and not affecting, in this aspect, the welfare of the animals.

### Verification of hematomas in the post-slaughter carcasses

In this study the presence and location of the hematomas were evaluated in the beef carcasses after skinning and the results are presented in Table 2.

Among the 33 cattle observed, only 5 animals did not present hematomas. Hematomas were observed in the regions of tail and croup, pelvic limb, flank, ribs and thoracic limb. The most affected region was the flank, followed by the croup and tail, as described in Table 2. The high percentage of contusions in these areas could be due to improper handling of the animals, the use of sharp objects, poles, electric shock or even trauma caused during the transportation from the rural property to the cold storage slaughterhouse, as described by Huertas et al. [33].

The pelvic limb represented the 3^rd^ most affected area. This high percentage was observed due to the lesions at the metatarsus (commercialized as hindshank muscle) of a certain batch which presented straight line marks that resembled the shape of a string. Therefore, it is assumed that these hematomas occurred due to the improper restraining at some time during the transportation. The route and the condition of the vehicle should also be considered, because poor road conditions and occupancy could lead to various contusions [31]. The present study did not identify cattle with severe wounds or deep hematomas and none of the animals died in the course of transportation.

**Table 2.** Hematomas detected in the beef carcasses divided according to the affected region in the carcass, number of animals affected and the percentage of animals with lesions in cold storage slaughterhouses from the Federal District and Surroundings.

| Region | Mean of the values obtained in the collections |  |  |  | Number of affected animals / total number of animals | Percentage of animals with lesions (%) |
| --- | --- | --- | --- | --- | --- | --- |
|  | Collection 1<br>n=9 | Collection 2<br>n=4 | Collection 3<br>n=7 | Collection 4<br>n=13 |  |  |
| Tail and croup | 2 | 1 | 5 | 9 | 17 / 33 | 51.5 |
| Pelvic limb | 5 | 2 | 0 | 0 | 7 / 33 | 21.2 |
| Flank | 5 | 2 | 3 | 7 | 17 / 33 | 51.5 |
| Ribs | 1 | 0 | 1 | 1 | 3 / 33 | 9.0 |
| Thoracic limb | 0 | 0 | 1 | 5 | 6 / 33 | 18.1 |
| Cervical region | 0 | 0 | 0 | 0 | 0 / 33 | 0.0 |
| Without hematomas | 3 | 0 | 1 | 2 | 5/33 | 15.1 |
| Total | 16 | 5 | 11 | 24 | 33/33 | - |

In the present study 5 animals did not present any type of hematoma, a value that corresponds to 15% of the animals. This is a positive result when compared to a study by Andrade et al. [35], which evaluated hematomas in cattle transported by river in the region of the Pantanal of Mato Grosso do Sul and found 5 animals without lesions among 88 evaluated, representing only 5.8% of the herd.

The combination of the results from the present study identified 11 animals with up to 1 hematoma, 8 animals with 2 and 14 with 3 or more hematoma lesions, results similar to those observed by Bertoloni et al. [34], where 60% of the slaughtered animals presented at least one hematoma in their carcass.

Among the most affected areas by hematomas, the hindquarter region was in second place in the second place, with results like Andrade et al. [35], who formulated the hypothesis of improper handling during the transportation and handling of the animals, which may have included the use of poles or electric stimuli to move them.

In addition to being used as a welfare indicator, hematomas generate economic loss to the producer, given that the meat can be compromised and discarded due to the lesion extension and depth [34]. Handling best practice can avoid great economic losses given that the main noble cuts are located in the most affected regions.

### Beef meat quality

The results of the pH analyses, shearing force test, cooking and drip weight losses and colorimetry of the meat samples obtained after 24h of cooling in the cold chambers of the industries are presented in Table 3.

A mean pH of 5.86 was observed in the animals of the present study. This pH value meets the parameter of normal meat (pH<6.0) [36]. The results also corroborate with Kuss et al. [5], in the state of Paraná, who evaluated young cattle (22 months old) entire with the mean 24h post-mortem pH of 5.9. These results are also similar to those observed by Andrade et al. [4] in cattle of the nelore breed, which found pH from 5.4 to 5.8, and to those in Batista de Deus et al. [28], in the state of Rio Grande do Sul, with the Aberdeen Angus breed, which identified values ranging from 5.6 to 5.78.

**Table 3.** pH, shearing force, cooking losses, drip losses and colorimetric (L*a*b*) analyses of the 33 beef meat samples (*Longissimus dorsi*) obtained after 24h in cold chamber of cold storage slaughterhouses located in the DF and Surroundings.

|  | Mean values obtained from the collections |  |  |  | Final mean<br>of all the<br>collections | Standard<br>deviation of<br>all the<br>collections |
| --- | --- | --- | --- | --- | --- | --- |
| Quality characteristics of<br>the meat samples | **1**<br>**n=9**<br>**Males 9**<br>**Females 0** | **2**<br>**n=4**<br>**Males 4**<br>**Females 0** | **3**<br>**n=7**<br>**Males 1**<br>**Females 6** | **4**<br>**n=13**<br>**Males 0**<br>**Females 13** | **1 – 4**<br>**n=33**<br>**Males 14**<br>**Females 19** | **1 – 4**<br>**n=33**<br>**Males 14**<br>**Females 19** |
| pH | 5.94 | 5.84 | 5.83 | 5.83 | 5.86 | 0.285 |
| Shearing force kg/f | 2.86 | 4.18 | 2.09 | 1.47 | 2.30 | 1.356 |
| Cooking losses % | 14.76 | 15.73 | 10.58 | 9.07 | 11.75 | 4.563 |
| Drip losses % | 0.70 | 1.46 | 1.70 | 2.93 | 1.88 | 1.354 |
| L* | 32.94 | 32.36 | 27.85 | 26.71 | 29.34 | 4.077 |
| a* | 14.35 | 15.25 | 13.34 | 14.82 | 14.43 | 2.526 |
| b* | 6.33 | 6.67 | 5.15 | 5.46 | 5.78 | 1.312 |

L\* - lightness; a\* - red color intensity; b\* - yellow color intensity

Only collection 01(with a pH value of 5.94) had a pH value approaching the high range (pH ≥ 6.0 according to Adzitey and Nuru) [27]. In the high range meat may present as DFD (dark, firm, dry). This may be associated with animals who have suffered a period of chronic or prolonged stress. This generates glycogen reserve in the muscle at the moment of the slaughter, leading to slow glycolysis with little lactic acid formation of and consequently a high final pH [32]. However, considering the results of this present study it is not possible to state that the observed pH in this collection is linked to a long travel time (prolonged chronic stress), given that the mean travel time was 1h 20m, a time shorter than the trips considered long (longer than 3h) [28].

No information was obtained on whether or not animals fasted in the facilities of origin, therefore, further studies are necessary to better characterize the pH of the region of the Federal District and Surroundings.

According to the statistical analysis of the relationship of pH with hematoma (presence and absence) in the beef carcasses of the present study, according to the Kolmogorov-Smirnov test, STATA 12®, there was no significant difference (p>0.05) between the means of the two variables. However, the same test highlighted a large difference in the standard deviation. This information indicated a trend where the pH of animals without hematoma lesions presented low variation among the groups (approximately 5.7 to 5.8), while the animals with hematomas presented a wide pH variation range (6.63 – 5.66), as presented next (Fig.1).

The tenderness test indicated an overall mean shearing force of 2.30 (kg/f) of all the samples, similar to that observed by Rubiano et al. [37] in Botucatu (SP), who observed mean values of 2.48 for cattle of the Canchim breed and 2.69 for Nelores. Fernandes et al., [38] reported for the same breed a mean value similar to that of the present study, of 3.09 in neutered males. These results accord with Lawrie [39], where values below 5 kg/f characterize the meat as tender.

Many factors can influence meat tenderness, such as the genetics, breed, gender, age at slaughter, diet, post-slaughter stress and cooling of the carcass [2]. The zebu breeds form most of the national herd, as they are better adapted to the challenges presented by the climate and parasites of the country [40]. However, the meat of these animals is considered tougher when compared to the bull breeds, the result of higher calpastatin activity in the zebus, an enzyme that inhibits calpain, which is responsible for tenderness [12].

**Figure 1.**
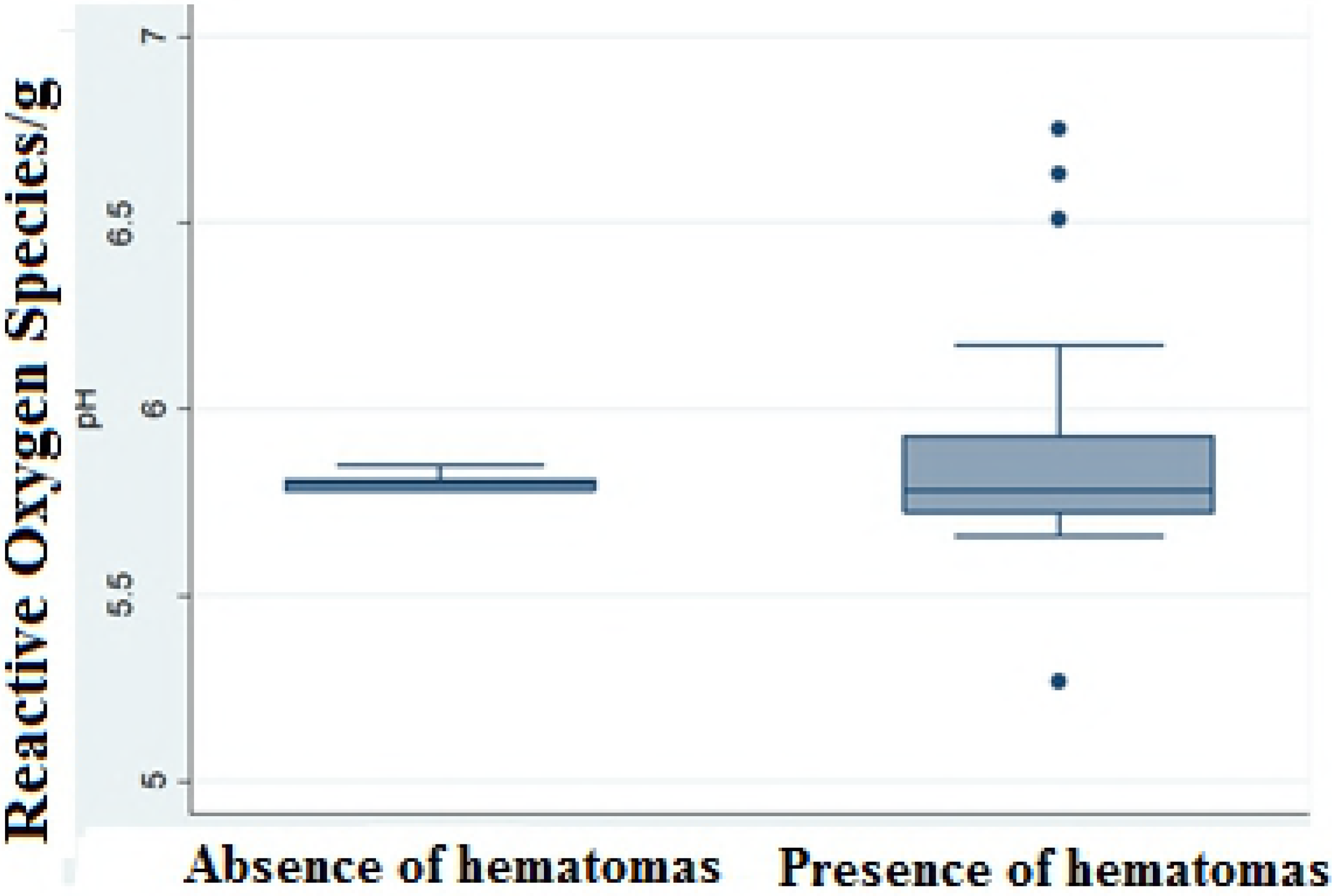
Statistical results presented in Boxplot according to the Kolmogorov-Smirnov Test, STATA 12®, for the evaluation of the relationship between the variables pH versus absence and presence of hematomas.

The present study indicated that all the animals were crossbred, presenting external characteristics of nelores and aged between 24 and 36 months old, however, it is not known what crossbreeding originated these animals. The small values presented in the shearing test may have been influenced by the crossbreeding as well as the presented sexual condition (females) that influence the age of slaughter, precociality and tenderness [5]. The present study did not identify neutered males with ages outside the range of 24 to 36 months old.

The statistical analysis performed by the Kolmogorov-Smirnov Test, STATA 12®, for the variables of gender (males and females) and shearing force for tenderness, identified a statistically significant difference between the means. The males presented a mean of 3.127 with standard deviation of 1.654, while the females presented a mean of 1.703 with deviation of 0.621, which leads to the conclusion that gender influences the tenderness of beef meat [12, 41].

When evaluated, the results of the present study confirm that the collections that were largely formed by females (collection 3 and collection 4) presented lower shearing force in comparison to the other groups, as previously demonstrated in Table 3.

According to the cooking loss test analysis, the present study found values ranging from 4 to 24.07% of total moisture loss, with an overall mean of 11.75%. Differently from the results obtained in this study, greater losses were observed by Barcellos et al. [42] in cattle of the nelore breed in Panama, with moisture losses of 24% and 29% in Angus x Nelore crossbreeds. Andrade et al. [4] found in cattle of the nelore breed greater losses of 29.1% when compared to the present study. Costa et al. [6], in Red Angus heifers losses between 20.1% and 25.5% and Menezes et al. (2005) [43] reported losses of 22.2% in Charolais and 22% in Nelores.

The coloration presented by the 33 cuts of approximately 500 grams obtained after 24h of cooling in cold chamber indicated mean values of 29.34 for L* (lightness), 14.43 for a* (intensity of the red color) and 5.78 for b* (intensity of the yellow color). The animals presented a slightly low L* value. According to Muchenje et al. [44], it is considered normal that when the pH is approximately 5.7, the L* values range between 33.2 - 41, a* between 11.1 – 23.6 and b* between 6.1 – 11.3.

Especially in collections 03 and 04 the L* presented the lowest values (27.85 and 26.71, respectively). When evaluated, the pH of these collections revealed means of 5.83 in both collections. A pH higher than 6.0 could characterize a DFD meat and therefore lower lightness (dark meat) with smaller water retention losses (dry) [32]. When evaluated individually, overall, most of the darkest meat samples (L* < 33) presented a pH higher than 5.8 (S1 Fig.).

A smaller lightness value represents a darker meat, which may be associated with the presence of fat, given that animals with higher content present higher reflectance [45], or even the breed, since the animals with a predominance of Nelore in their genotype presented darker meat, a behavior related to the agitated temper of the breed [5], another factor that could be attributed to the pH in the case of meat with low lightness, because meats with values higher or equal to 6.0 tend to have low light reflectance due to high water retention capacity [47]. In turn the a* and b* values are in accordance with the previous studies, such as Barcellos et al. [42] for a* of 14.58 and Andrade et al. [4] for b* of 3.78, all performed with cattle of the Nelore breed.

A variety of other factors could influence the coloration, such as the activity level of the animal: pasture-raised animals exercise more and are slaughtered at a more advanced age, thus presenting higher myoglobin content and having a more intense red color [47].

Drip loss is one of the parameters used to evaluate the water retention capacity (WRC). In the present study, the mean of the collections for the drip losses was 1.88%, unlike those observed by Igarassi et al. [47], who found the value of 4.30% for the drip losses in Red Angus x Nelore animals in Botucatu (SP). Strydom et al. (2011) [49] found the value of 2.01% in cattle of the Brahman breed, Hopkins et al. [50] found a mean of 2% in 115 carcasses from cold storage slaughterhouses and Lage et al. [51] found the value of 1.85% in Nelores.

The present study indicated low fluid loss, once again, due to the high water retention capacity, a result attributed to the low protein denaturation and high water bond, as described by Adzitey and Nurul [36].

### Means of electron paramagnetic resonance in the samples of muscle tissue

The detection of the reactive oxygen species in the samples of muscle tissue collected post-slaughter was performed using a Bruker® EMXplus EPR X-Band spectrometer. The reading of each sample generated a spectrum where the peak-to-peak amplitude of the central transition is proportional to the concentration of unpaired electron spins, from the molecules of the spin markers that produce a signal with amplitude amplified by a Lock-in, an indicator of the number of free radicals present per gram of sample. After converting the values of the curve in the proper equation, a numeric value is obtained, which represents the amount of reactive oxygen species detected per gram of tissue. The mean values obtained for the samples of each collection period are found in Table 4.

**Table 4.**
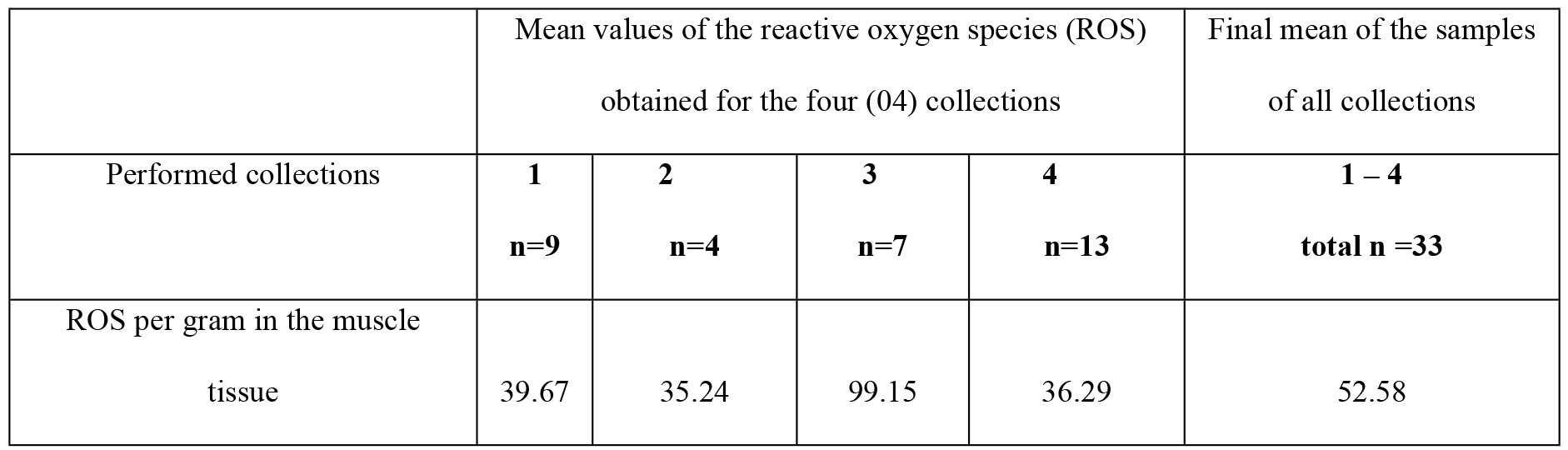
Mean of the values for the reactive oxygen species obtained by the EPR curve for the muscle tissue samples, per gram.

There are no results of ROS quantification in muscle tissue obtained in post-slaughter of beef meat. Gadjeva et al. [17] used the EPR technique for the detection of free radicals and antioxidant agents in different muscle groups in pork meat *in natura,* in the range of 2 to 3 hours after slaughter. The researchers detected in the *Longissimus* muscle a mean value of 7.69 ±0.91 of ROS per gram, a lower value compared to the means of the other muscles analyzed in the study, the subscapularis (8.66 ±1.17) and gluteobiceps muscles (8.54 ±1.05). In the present study the analyzed muscle piece was the extensor carpi muscle and a higher value was detected in the samples than that found by Gadjeva et al. [17], with the mean value of 52.58 ROS/g for all the collections.

The implications of this high mean are not clear because there still is a lack of studies on the ROS dosage in meats of different domestic species. Hence, some hypotheses can be formulated, which would explain these discrepant results. Firstly, the different species evaluated, since the present study performed the detection of ROS in beef meat samples, while Gadjeva et al. [17] worked with pigs, so it is not possible to compare the results. Other factors that may have influenced the amount of ROS are the selected muscle region and the applied methodology. In this studythe extensor carpi muscle piece was analyzed, collected immediately after slaughter and frozen in liquid nitrogen, while Gadjeva et al. [17] samples were collected from three other muscle regions – pieces from the longissimus, subscapularis, gluteobiceps muscles – removed 3 hours after slaughter on average. which would mean that antioxidant enzymes had been acting for 3h.

Gadjeva et al. [17] present evidence that the Catalase and Superoxide dismutase enzymes decrease the content of free radicals in the muscle cell, decreasing the local oxidative activity in the range of 2 to 3 hours post mortem, since it was observed that when oxidative stress increases, the activity of these enzymes also increases, demonstrating the response of the antioxidant enzymatic system in fresh pork meat. Because in the present study the samples were frozen, these enzymes probably did not have time to act This is one of the most influential surveyed factors for the large number of free radicals detected by the EPR in this study. Therefore, it is necessary to perform further studies and to quantify the reactive oxygen species with different collection and storage conditions and post-slaughter time period.

Further studies should be performed on this subject, comparing different protocols for the measurement of free radicals with the qualitative analysis of the meat, so it is possible to verify whether the presented amount of ROS is enough to affect the product quality.

The analysis of the variables ROS in the tissue and hematomas (absence or presence) according to the Kolmogorov-Smirnov Test, STATA 12®, indicated no significant difference in the averages evaluated, while there was a difference in the standard deviation results, revealing a trend where animals without hematomas have a smaller and narrower variation rate when compared to animals that presented hematomas (Fig. 2).

**Figure 2.**
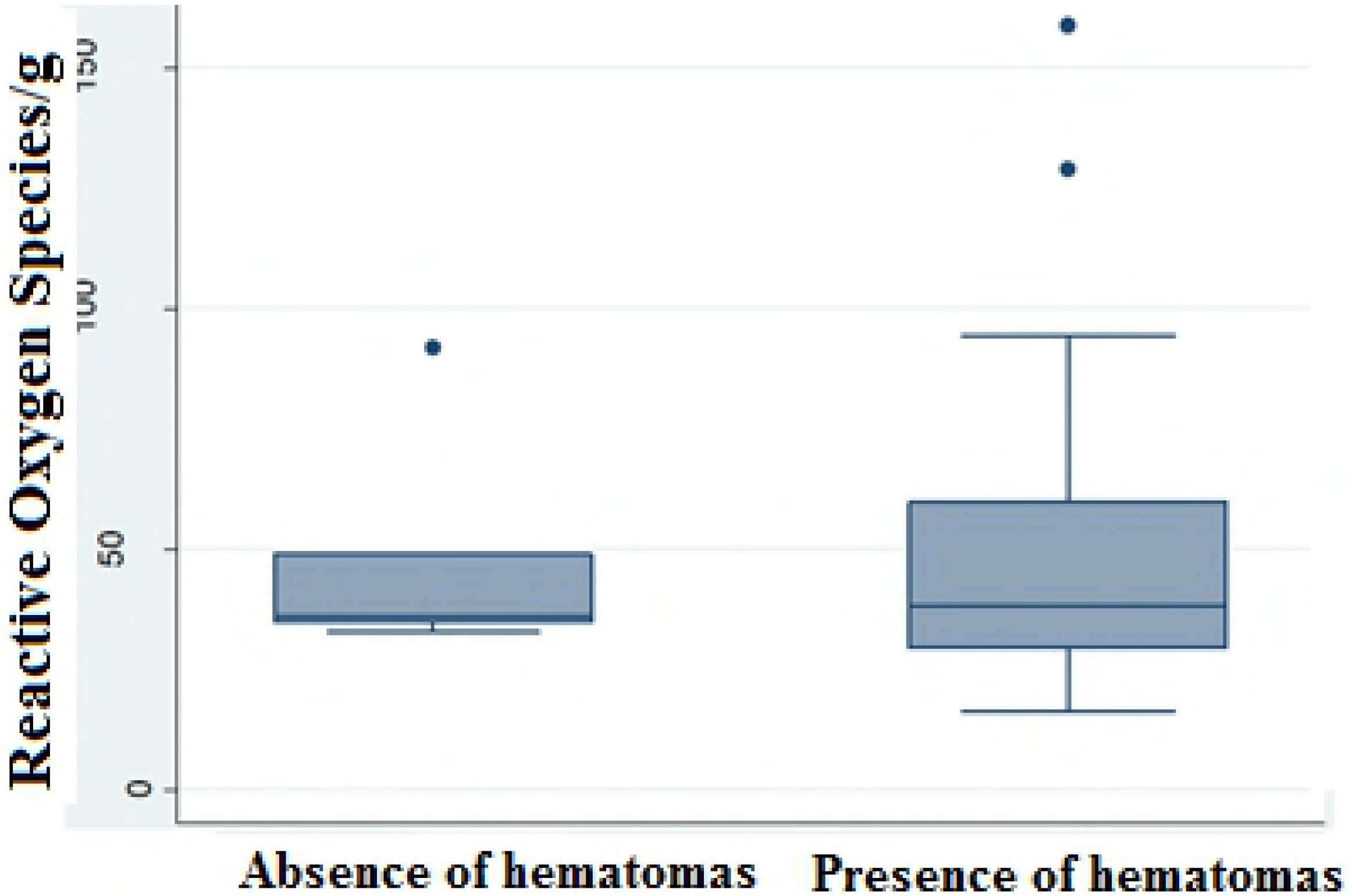
Results presented in Boxplot by the Kolmogorov-Smirnov Test, STATA 12®, for evaluation of the relationship between the ROS variables in the tissue/gram versus the absence and presence of hematomas.

Although many studies analyze beef meat quality, most are performed comparing meat obtained from different breeds, genetic crossbreeding types and different diets [4, 5, 6, 7]. The present study is the first to characterize the quality of beef meat from cold storage slaughterhouses in the region of the Federal District and Surroundings, and, overall, the results met the desirable quality parameters.

Studies that evaluate the presence of free radicals in food such as meat are more easily found, however, most of them focus their study on the detection of free radicals in products that underwent some treatment with an antioxidant action [52, 53, 54].

The present study is the first performed in Brazil that relates pre-slaughter cattle stress with the detection of ROS, and its influence on meat quality. However, it is necessary to perform further studies to allow technical standardization, more clarity in the reading of results and the verification of a possible relationship between the factors of pre-slaughter stress, the presence of free radicals and meat quality.

## Conclusion

According to the results of the present study, the distance and time taken by the animals to arrive at the cold storage slaughterhouses in the region of the Federal District and Surroundings and the rest time during which they remained in the slaughterhouse were not enough to influence their meat quality.

The results obtained in the organoleptic analyses of the meat indicated some variations in the evaluated parameters, however, the samples met the parameters considered for high-quality meat. Statistically, two trends were observed in the present study. According to the Kolmogorov-Smirnov, test analysis for the evaluation of the relationship between the variables of pH versus the absence or presence of hematoma, it was proved that the pH value can be affected by the presence of hematomas. Therefore, the results presented by the Kolmogorov-Smirnov test for the assessment of the relationship between the ROS variables in the tissue/gram versus absence and presence of hematomas, which demonstrated that the amount of reactive oxygen species in the muscle tissue can be influenced by tissues with hematomas.

No studies were found comparing the amount of pre-slaughter ROS in relation to stress and beef meat quality, thus requiring further studies in this area to verify whether there is a possible relationship between these parameters and whether they are correlated to stress.

## Acknowledgements

To the Coordination for the Improvement of Higher Education Personnel (CAPES) for granting the two-year masters scholarship, to the University of Brasília for providing this research opportunity, to my advisor and professor Dr. Angela Patricia Santana and to the project co-advisor Paulo Eduardo Narciso de Souza and to professor Dr. Fabiane Hiratsuka Veiga de Souza. I also thank for the assistance provided, the following laboratories UnB: Laboratory of Food Microbiology. School of Agronomy and Veterinary Medicine, Laboratory of Electron Paramagnetic Resonance. Institute of Physics, Laboratory of Biochemistry and Protein Chemistry. Institute of Biological Sciences and Laboratory of Veterinary Epidemiology. School of Agronomy and Veterinary Medicine.

## Support Information

**Data table with all the performed collections**

